# Buffer Valency Engineering Enables High-concentration and Shelf-stable DNA Transfection Particles for Viral Vector Production

**DOI:** 10.1101/2025.07.02.662322

**Authors:** Jinghan Lin, Yizong Hu, Turash H. Pial, Kailei D. Goodier, Di Yu, Marine Guise, Paetra Brailsford, Maria Choi-Ali, Sixuan Li, Yining Zhu, Jingyao Ma, Leonardo Cheng, Xiaoya Lu, Nicole Korinetz, Tza-Huei Wang, Tine Curk, Hai-Quan Mao

**Author notes:** Present address: David H. Koch Institute for Integrative Cancer Research, Massachusetts Institute of Technology, Cambridge, MA, USA. These authors contributed equally to this work (Jinghan Lin, Yizong Hu). Corresponding authors: Yizong Hu, and Hai-Quan Mao.

## Abstract

Cost-effective and scalable production is critical for advancing the clinical translation of adeno-associated virus (AAV)-mediated gene therapy. The widely used transient transfection method using plasmid DNA (pDNA)-loaded transfection particles for AAV production faces technical challenges due to instability of the particles and the concentration limits for particle preparation, hindering reproducibility and scalability. Here, we report a streamlined and scalable strategy to generate shelf-stable, highly concentrated pDNA/poly(ethylenimine) (PEI) transfection particles. By incorporating trivalent citrate ions in the dilution buffers, we kinetically modulate electrostatic complexation to achieve uniform nanoparticle assembly and prevent aggregation at high concentrations. This enables a tenfold increase in pDNA concentration in stabilized transfection particles from a typical range of 10–20 μg/mL to 200 μg/mL, while reducing the required dosing volume from 5–10% to 0.5% of the cell culture medium. The particle assembly process is robust to changes in mixing scale and timing and is compatible with standard workflows. We demonstrate equivalent AAV production efficiencies to standard methods and consistent performance in various production scales, which confirms the practical utility of this assembly method in developing robust, scalable, and cost-effective AAV manufacturing processes.

## Introduction

Gene therapy has evolved as a transformative modality for treating disease and improving healthcare, using both non-viral carriers^1^, such as lipid^2^ and polymer^3^-based nanoparticles, and viral vectors including lentivirus, adenovirus, and adeno-associated virus (AAV). Since its discovery in 1965, AAV has become a leading platform due to its high transduction efficiency in dividing and non-dividing cells, prolonged transgene expression, and low pathogenicity, as demonstrated in clinical trials^4–6^. AAVs are now the most widely used viral vectors in gene therapy, prompting significant efforts to optimize their large-scale production for clinical applications^7–11^. However, persistent challenges in vector packaging and purification continue to hinder cost-effective manufacturing, which is an essential step toward making these therapies more accessible and broadly applicable.

The industry standard method for AAV production involves co-transfection of production cell lines, such as human embryonic kidney 293 (HEK293) cells, with transfection particles complexed by polyethyleneimine (PEI)^12^ and a cocktail of three plasmid DNAs (pDNAs) encoding AAV assembly components^11, 13, 14^. While this method has been refined for more than two decades, it is plagued by limitations associated with the pDNA/PEI transfection particles, such as a lack of understanding of critical properties that govern transfection efficiency^15, 16^, batch-to-batch variability^17^, poor stability^18, 19^, and complexation behaviors that are inconsistent across different scales and processes^20^.

Previously, we identified the colloidal size of pDNA/PEI transfection particles (using PEIpro^®^ from Polyplus Sartorius) as a critical parameter determining their transfection efficiency, with an optimal size of 400 to 500 nm^21^. We then developed a stepwise, electrostatic charge-mediated assembly method that enabled the manufacturing of bench stable (> 4 h at room temperature) and shelf-stable (> 1 year at –80°C) 400-nm pDNA/PEI particles and successfully benchmarked their performance in producing lentiviral vectors^21^. With specifically designed flow mixing devices and protocols that enabled continuous-flow preparation of pDNA/PEI particles to improve the reproducibility of cell transfection, the assembly method was proven scalable to a commercially relevant scale, producing such transfection particles at a flow rate of 1 L/min^22^. Despite these improvements, the assembly method requires specialized pumping and turbulent mixing equipment, a dedicated particle manufacturing pipeline, and a time-consuming manufacturing process^22^, which are sub-optimal for adoption in large-scale bioreactors.

Here, we present a simplified method to prepare shelf-stable pDNA/PEI transfection particles based on kinetic control of the pDNA/PEI polyelectrolyte complexation (PECn) process. By introducing multivalent counterions to PEI, specifically trivalent citrate ions, we reduced the rate of PECn, such that the mixing rate of the pDNA and PEI components is less relevant. This kinetic relationship permits further assembly and growth of pDNA/PEI particles from uniform seed complexes, even at an exceptionally high pDNA concentration. Compared to the original phosphate buffer, citrate ions mediate significantly more uniform assembly during the particle growth process. This more controlled process allows the preparation of particles using a variety of mixing techniques with a high degree of consistency across different preparation scales.

Using citrate-mediated assembly, we successfully validated a protocol to produce shelf-stable pDNA/PEI transfection particles using PEIpro at 200 μg pDNA/mL, producing transfection particles at 10 to 20 times higher concentration than industry practice, which can correspondingly reduce particle dosing volume by an order of magnitude in bioreactors. We also compared the performance of these shelf-stable transfection particles in AAV production with those prepared with industry-developed processes in collaboration with two independent commercial laboratories.

## Results and Discussion

### Kinetics analysis of the manufacturing process for pDNA/PEI transfection particles

The assembly of pDNA/PEI particles with a target size range of 400–500 nm involves two stages (**Fig. 1a**): the initial polyelectrolyte complexation (PECn) and subsequent nanoparticle assembly (NPa). During PECn, the mixing rate of pDNA solution and PEI solution compared to the complexation rate of the two macromolecules is critical for the ability to generate uniform polyelectrolyte complex (PEC) nanoparticles. The PECn “reaction” is highly energetically favorable; thus, the characteristic complexation time τ_e_, *i.e.*, the time it takes to complete the complexation between pDNA and PEI, is typically shorter than tens of milliseconds^23^. On the other hand, when pDNA and PEI solutions are mixed using conventional methods, the relatively low and mismatched diffusivities of the two types of macromolecules, due to the large difference in their hydrodynamic size and molecular weight, make homogeneous mixing difficult to attain. Importantly, the theoretical characteristic mixing time τ_m_, *i.e.*, the time it takes to achieve homogenous mixing of pDNA and PEI components, is two to three orders of magnitude longer than τ_e_, yielding PEC nanoparticles with a higher degree of size heterogeneity^23^.

**Figure 1.**
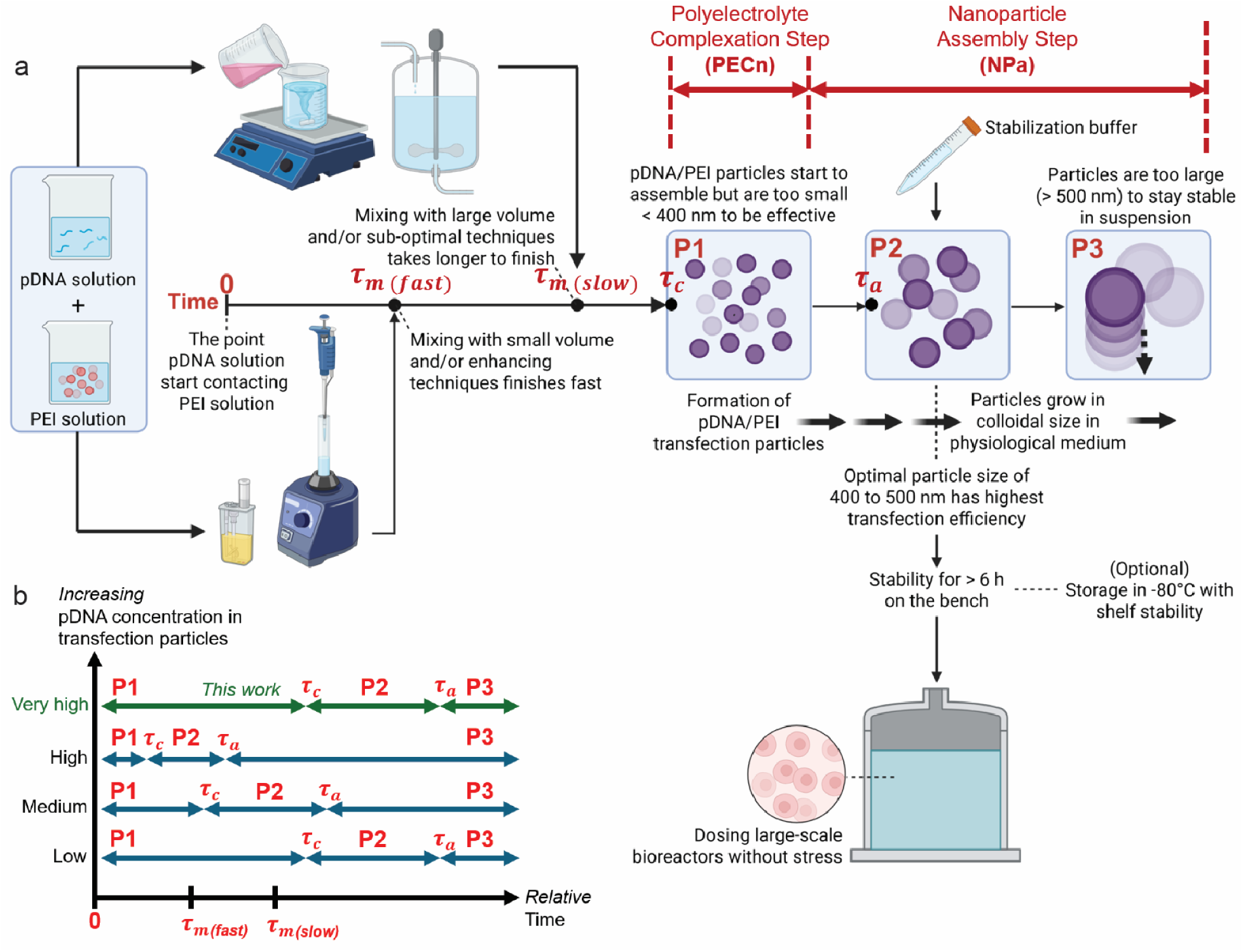
Kinetically controlled assembly process of the shelf-stable pDNA/PEI transfection particles. **(a)** Schematic overview of the process to prepare pDNA/PEI transfection particles, involving mixing of pDNA cocktail solution and PEI solution that is followed by polyelectrolyte complexation (PECn) and the nanoparticle assembly (NPa) in a medium that bears physiological pH and ionic strength. The size growth of pDNA/PEI particles can be divided into three periods: P1, in which particles possess a size < 400 nm; P2, in which particles possess a size between 400 and 500 nm; P3, in which particles possess a size > 500 nm. **(b)** Comparisons of the relative time scales of the mixing, PECn, and NPa steps. The figure was generated by BioRender.com.

One approach to address this challenge is to speed up the mixing rate and mixing efficiency, thus reducing τ_m_ by reducing the diffusion distance of the two components^24, 25^. Previously, we adopted a flash nanocomplexation (FNC) technique^23^ to break the two solution jets into micron-scale sheets through a turbulent mixing regime in a microchamber, drastically reducing the diffusion distance between the two components. Under this condition, all pDNA molecules and PEI molecules complex nearly simultaneously, thereby generating uniform PEC nanoparticles with a high density of residual positive charges that render them colloidally stable. These PEC nanoparticles are then aggregated under controlled conditions to obtain the 400–500 nm transfection particles. However, the implementation of such advanced mixing technology and multi-step procedures may not always be favorable in the development of scale-up transfection processes in bioreactors.

Here, we propose a new streamlined method to substantially slow down the rate of PECn, thus increasing τ_e_ to match the slower mixing process, thereby improving the uniformity of the PECn. We hypothesize that using the citrate ion as a multivalent and reversible “chelator” to PEI can serve this purpose. Citrates have a small molecular weight and a much higher diffusion rate to “pre”-complex with PEI during any mixing process. They can reversibly reduce the density of positive charges on PEI, which can markedly reduce the energetic driving force and complexation rate with pDNA. This becomes particularly important during PECn step with a higher pDNA concentration with an even shorter τ_e_ as determined by reaction kinetics (**Fig. 1b**). Due to the higher negative charge density on pDNA, the pDNA molecules are thermodynamically favorable to form PEC with PEI, such that the citrate ions are eventually released from the PEC. We anticipate that this method of generating PEC nanoparticles will be less restricted by the mixing rate of the components, *i.e.*, τ_e_ ≥ τ_m_ (**Fig. 1b**). This kinetic control enables more uniform PEC formation regardless of mixing method or scale.

Following mixing and the initial PECn process, the surface charges of the pDNA/PEI nanoparticles are drastically reduced due to charge neutralization, and the charge screening conferred by the citrate ions and the high ionic strength in the buffer reduces the barriers of assembly and permits nanoparticles to “aggregate” into larger particles. This NPa process is a relatively slow “reaction,” with a longer characteristic time of τ_a_, defined as the time it takes to reach the target size of the particles (**Fig. 1a**), due to the much slower diffusion rate of the nanoparticle units compared with the original pDNA and PEI macromolecules. During this NPa step, we hypothesize that pDNA/PEI particles can form more uniform aggregates at a controlled rate because of the more uniform PECn process mediated by citrate ions. Once the particles grow to the target size, a stabilization buffer dissolved in an acidic pH is used to effectively re-protonate the PEI molecules on the surface of the PEC particles, which can arrest the particle growth and preserve particle uniformity via long-range electrostatic repulsion^21, 23^.

### Citrate-buffered saline enhances uniformity and stability of high-concentration assembly of pDNA/PEI particles

In our previous method, the phosphate-buffered saline (PBS, pH 7.0) was used to induce particle growth^21^ by deprotonating nearly 60% of the amine groups on PEI^26^. The salts in the buffer, including Na^+^, K^+^, Cl^-^, and phosphates, provide charge screening to PECn. Here, we modified this buffer to replace a portion of the buffer salts with trisodium citrate, while retaining the same ionic strength, yielding citrate buffered saline (CBS). The pH of CBS was tuned to be 7.0. In all the experiments, we targeted a high pDNA concentration and selected the working concentrations of pDNA and PEI solutions to be 440 μg/mL and 580.8 μg/mL, respectively, correlating with a nitrogen-to-phosphate (N/P) ratio of 5.5. In an initial run, we used pipetting as the mixing method to add 100 μL of PEI solution into an equal volume of the pDNA working solution diluted in either PBS or CBS. Continuous DLS measurements revealed that the overall particle growth rates in both PBS and CBS buffers are comparable (**Fig. 2a, b**), and the average diameter of the particles reached the target size of 400 nm at around 20 min after mixing. However, a sharp difference in the polydispersity index (PDI), a measure of particle uniformity (see definition in **Methods**), was observed. The PDI increased with the average size of the particles assembled in PBS (**Fig. 2c**), suggesting increasing size heterogeneity as the particles grew, however, the PDI of the pDNA/PEI particles assembled in CBS remained largely unchanged in a range of 0.05 to 0.2 throughout the entire growth process, which is considered reasonably uniform^27^ (**Fig. 2d**).

**Figure 2.**
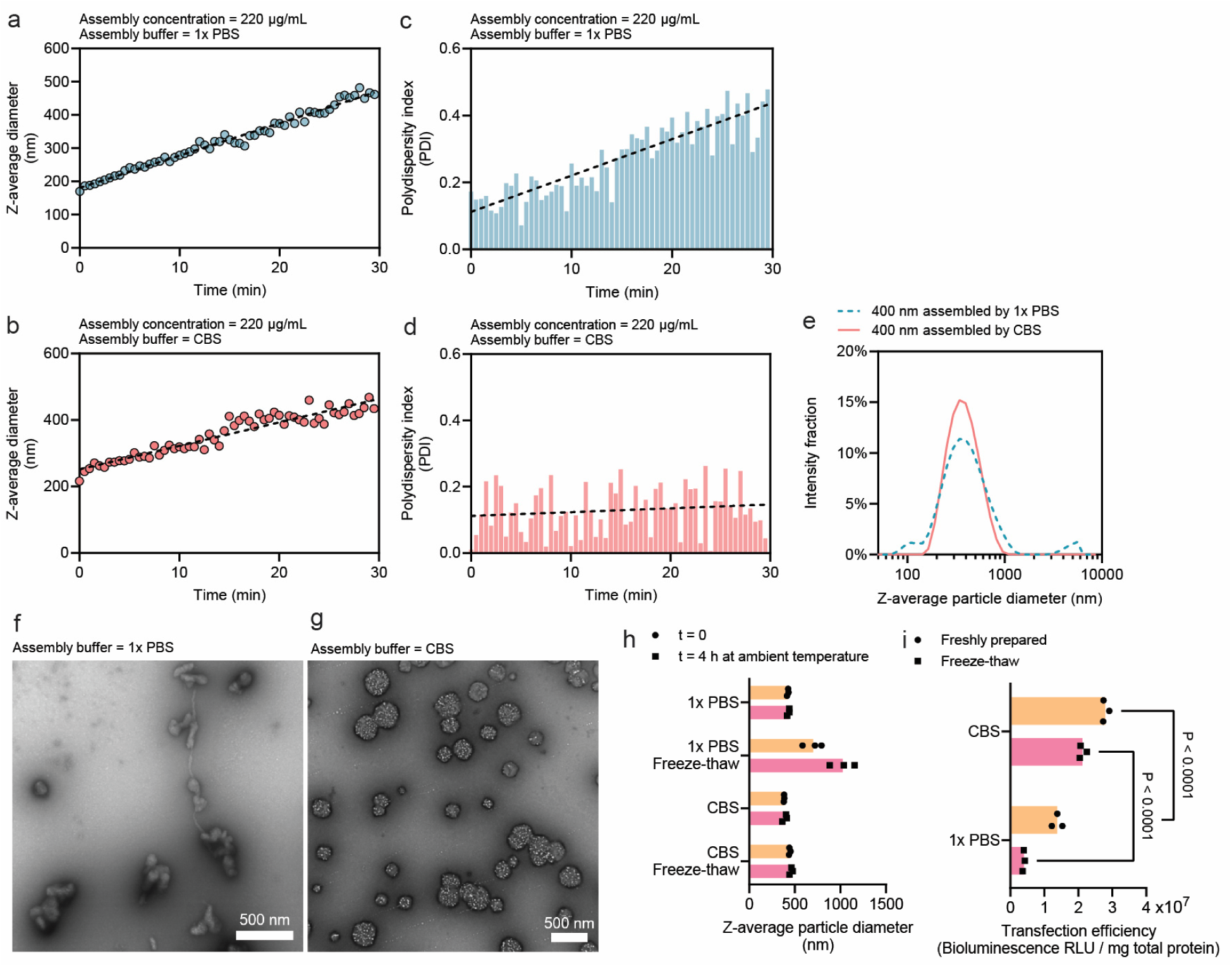
Citrate-mediated assembly of pDNA with PEI generates uniform and stable particles. (**a, b**) The size growth curve of the assembly of pDNA/PEI particles monitored by consecutive dynamic light scattering (DLS) measurements in the assembly buffer of 1× PBS **(a)** or citrate buffered saline, CBS **(b)**. **(c, d)** The polydispersity index (PDI) of the growing pDNA/PEI particles measured for the assembly buffer of 1× PBS **(c)** or CBS **(d)**. **(e)** The size distribution given by DLS of 400-nm particles assembled by either PBS or CBS buffer. **(f, g)** Transmission electron microscopy (TEM) assessing the uniformity of 400-nm particles assembled by 1× PBS **(f)** or CBS **(g)**. **(h, i)** The on-bench (ambient temperature) and freeze-thaw stability of 400-nm particles produced by either buffer in terms of the particle size assessed by DLS **(h)**, or the transfection efficiency on HEK293T cells **(i)** when the particles were loaded with 10% (w/w) luciferase pDNA.

Once the pDNA/PEI particles reach the target size of 400 nm, the particle growth can be arrested by the addition of a stabilization buffer, which contains hydrochloric acid to protonate surface PEI to the maximum degree. Trehalose (19%, w/w) was added to this stabilization buffer to confer stability during cryo-storage. The size stability was verified by DLS, showing a more uniform size distribution of the CBS-assembled particles (**Fig. 2e**), and by transmission electron microscopy (TEM), examining the morphology of the particles. The pDNA/PEI particles assembled by PBS appeared to be randomly shaped agglomerates with drastically different sizes (**Fig. 2f**), whereas the particles assembled by CBS are more uniform in size with a predominantly spherical shape (**Fig. 2g**). Both particles assembled in PBS and CBS buffers remained stable under the ambient temperature for at least 4 h; nonetheless, only particles assembled in CBS remained stable upon a freeze-thaw cycle (**Fig. 2h**). More importantly, when we encapsulated luciferase pDNA in the particles and transfected HEK293T cells using stabilized particles, 400-nm pDNA/PEI particles assembled in CBS showed a significantly higher transfection efficiency than those prepared in PBS (**Fig. 2i**), presumably due to the more uniform size distribution within the optimal size range of 400 to 500 nm.

Of important note, further increasing the PBS concentration, which theoretically increases the degree of charge screening at a higher ionic strength, did not provide a similar benefit to CBS-mediated assembly, but rather resulted in uncontrollable aggregation **(Supplementary Fig. S1**). We also examined the effect of citrate triphosphate concentration in the CBS buffer as multivalent ions on particle supramolecular assembly^28^, the NPa step between τ_e_and τ_a_, that particle growth could not occur when citrate concentration was too low, and uncontrollable aggregation occurs when citrate concentration was too high (**Supplementary Fig. S2a**). The detailed kinetics analysis of the effects of multi-valent ions on τ_a_, and the relationship between τ_e_and τ_a_, are beyond the scope of this manuscript and warrants further investigations. Nonetheless, the selected citrate concentration could accommodate a wide range of pDNA concentrations to mediate reproducible particle growth (**Supplementary Fig. S2b**).

### Molecular dynamics simulation reveals mechanistic insights into the kinetic modulation of the pDNA/PEI particle assembly by CBS

To explicitly examine the molecular complexation behaviors between pDNA and PEI in PBS and CBS, we conducted molecular dynamics (MD) simulations. We employed a coarse-grained model by condensing 3–4 heavy atoms (e.g., nitrogen, carbon, oxygen, phosphate) into a single particle^29^. This captures the necessary chemical characteristics of the repeating units on each macromolecule, while allowing simulation of sufficiently large length and time scales. The PEI was modeled as a linear polymer chain with 40% of its nitrogen atoms protonated, mimicking physiological conditions^26^.

Through DLS, we found that diluting PEI into PBS or CBS only induces a minimal degree of self-assembly of PEI with the counter ions, characterized by a slight increase in the diameter of PEI without further assembly (**Fig. 3a**). MD simulation showed a similar outcome that at equilibrium, PEI-phosphate interaction and PEI-citrate interaction were apparent (**Fig. 3b**) without a significant increase in their number-average cluster size over time (**Fig. 3c**). The fluctuations observed in the number-average cluster size of PEI-citrate shows that the citrate-PEI binding is reversible.

**Figure 3.**
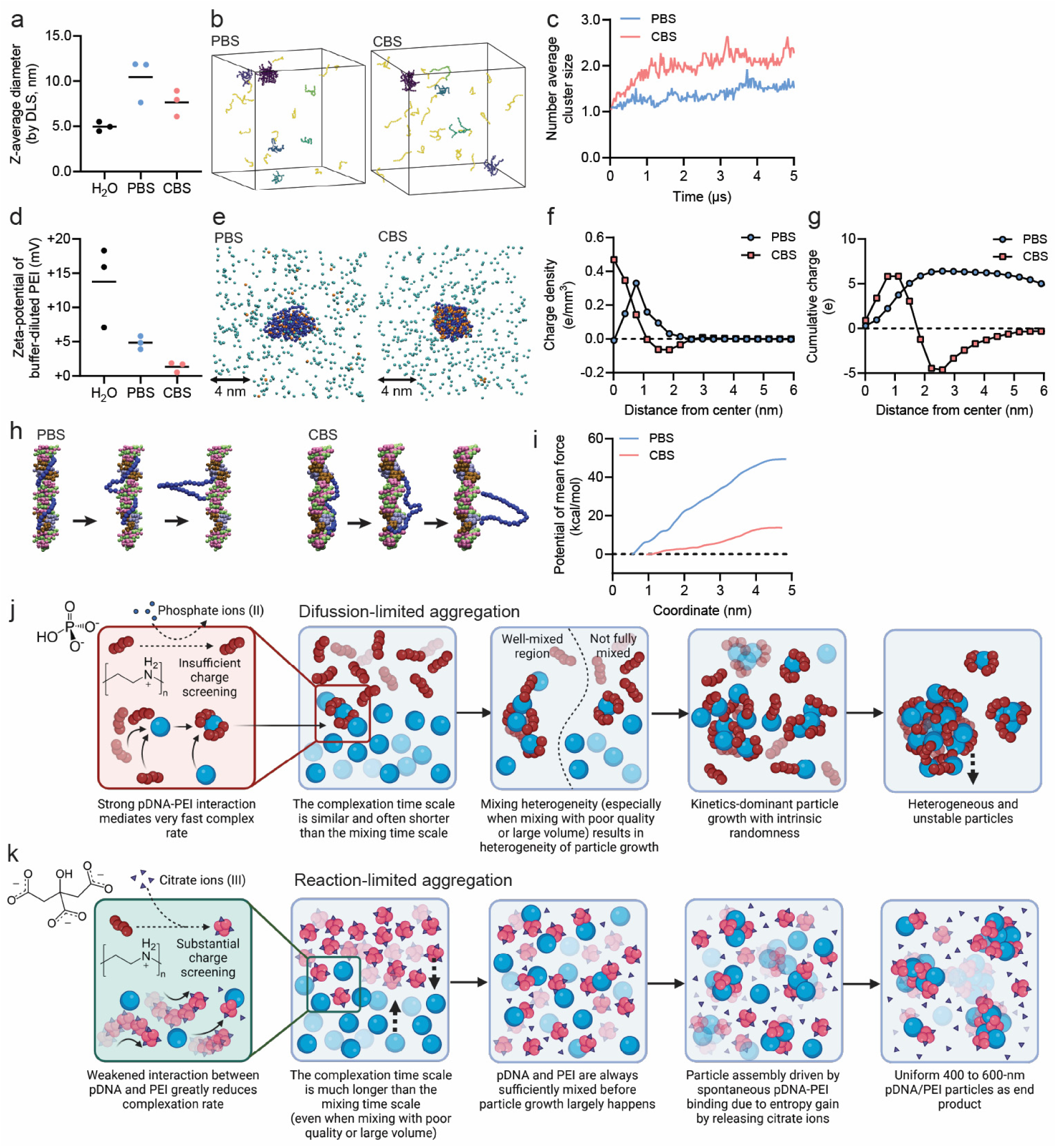
Mechanism understanding of pDNA/PEI particle assembly mediated by PBS vs. CBS. **(a)** The size of PEI molecules upon dilution in different buffers. **(b, c)** MD simulations **(b)** illustrating aggregation behavior of PEI molecules independent of pDNA in different buffers and quantifying the number average cluster size over time **(c)**. In **(b)**, different color indicates different clusters, yellow indicates unassociated single chains. **(d)** The zeta-potential of PEI molecules upon dilution in different buffers. **(e**–**g)** MD simulations **(e)** illustrating ion-PEI binding (Blue: PEI chain; Orange: multi-valent ions; Green: mono-valent ions) and quantifying **(f)** the charge density and **(e)** the cumulative charge distribution from the center of mass of PEI towards the surface. **(h, i)** MD simulations of DNA/PEI complexation in PBS and CBS. **(h)** Snapshots of umbrella sampling simulations along different pulling coordinates and **(i)** the free energy of pDNA/PEI binding in different buffers. **(j, k)** Schematic overview of the hypothetical mechanisms governing pDNA/PEI assembly in PBS **(j)** or CBS **(k)**.

Since complexation is driven by charge, we measured the zeta potential of PEI molecules by phase analysis light scattering (PALS). Substantial charge screening was observed in both CBS and PBS (**Fig. 3d**), with CBS yielding nearly full charge neutralization. To elucidate the molecular basis of the observed charge screening, we performed MD simulations resolving the spatial charge distribution around a single PEI chain with a realistic polymer size (19 kDa). The simulations revealed that the number of condensed ions is significantly higher in CBS than in PBS (**Fig. 3e**), indicating a weaker association between phosphate ions and PEI. The more diffuse ion cloud of PEI in PBS may also explain the larger hydrodynamic size of PEI molecules in PBS characterized by DLS (**Fig. 3a**), while the difference is marginal and within error bars. of This behavior can also be observed in radial charge density profile (**Fig. 3f**) where citrate ions assembled into a well-defined shell around the PEI chain, generating a net negative charge density near the polymer interface (**Fig. 3e, f**). As a result, citrate ions achieved near-complete charge compensation, leading to a cumulative charge approaching zero at long distances, consistent with the stronger charge neutralization observed in PALS measurements (**Fig. 3d**). In contrast, phosphate ions failed to fully neutralize the positive charges on PEI, reflected in a net positive cumulative charge profile (**Fig. 3g**) and non-zero zeta-potential in PALS measurements (**Fig. 3d**).

Further MD simulations in PBS and CBS were performed to elucidate the effect of divalent and trivalent ions on DNA–PEI interactions. A single PEI molecule (degree of polymerization = 28, ∼1.2 kDa) was placed in a simulation box containing a 24-base-pair DNA duplex and PEI–DNA binding free energy was quantified in both buffers using umbrella sampling (**Fig. 3h**). The resulting profiles (**Fig. 3i**) show that binding is markedly weaker in CBS than in PBS, demonstrating that citrate ions significantly attenuate PEI–DNA affinity.

Strong PEI–DNA interactions in PBS impede structural rearrangement, leading to kinetically trapped, irregular complexes; characteristic of a **diffusion-limited aggregation** mechanism. In contrast, weaker interactions in CBS allow extensive structural reorganization, facilitating the formation of more spherical complexes. Here complex growth is governed by the slower association/reorganization or ion replacement step, i.e., **reaction-limited aggregation** in the citrate-mediated system. Even though PEI carries near-zero net charge in CBS, binding remains spontaneous because displacement of bound citrate ions into bulk solution provides an entropic driving force. Collectively, these results suggest that PBS-mediated assembly proceeds rapidly (short τ_e_) and is prone to heterogeneity, whereas CBS-mediated assembly is slower (longer τ_e_) and yields more homogeneous complexes, largely independent of mixing rate (**Fig. 3j,k**). In further self-assembly simulations (**Supplementary Fig. S3**), we successfully captured the morphology differences of DNA-PEI complex depending on charge neutralization by ions, matching the experimental observations (**Fig. 2e–g**).

### Robustness of the kinetic assembly in CBS under different mixing conditions and scales

Next, we tested the hypothesis that the slower PECn kinetics rendered by citrate ions can ensure uniform assembly under different mixing conditions. To quantitatively vary the mixing quality, we used a confined impinging jet (CIJ) mixer^30^ coupled with digital syringe pumps to control the flow rate during mixing. With a defined geometry (**Fig. 4a**), we previously demonstrated that the characteristic mixing time τ_m_correlates with the total flow rate (**Supplementary Table 1**)^23, 31^. We varied the total flow rate to mix PEIpro solution with pDNA solution diluted by CBS to obtain a characteristic mixing time ranging from 15 milliseconds to minutes, pipetted the stabilization buffer into the mixture at the same post-mixing time, and finally vortexed the mixture to fully stabilize the particles. Using this experimental set-up to assemble the 400-nm pDNA/PEIpro particles by CBS, the PDI of the particles remained close to 0.1 regardless of the mixing rate (**Fig. 4b**). All particles generated showed consistent transfection efficiency using luciferase pDNA as a reporter to transfect HEK293T cells (**Fig. 4c**).

**Figure 4.**
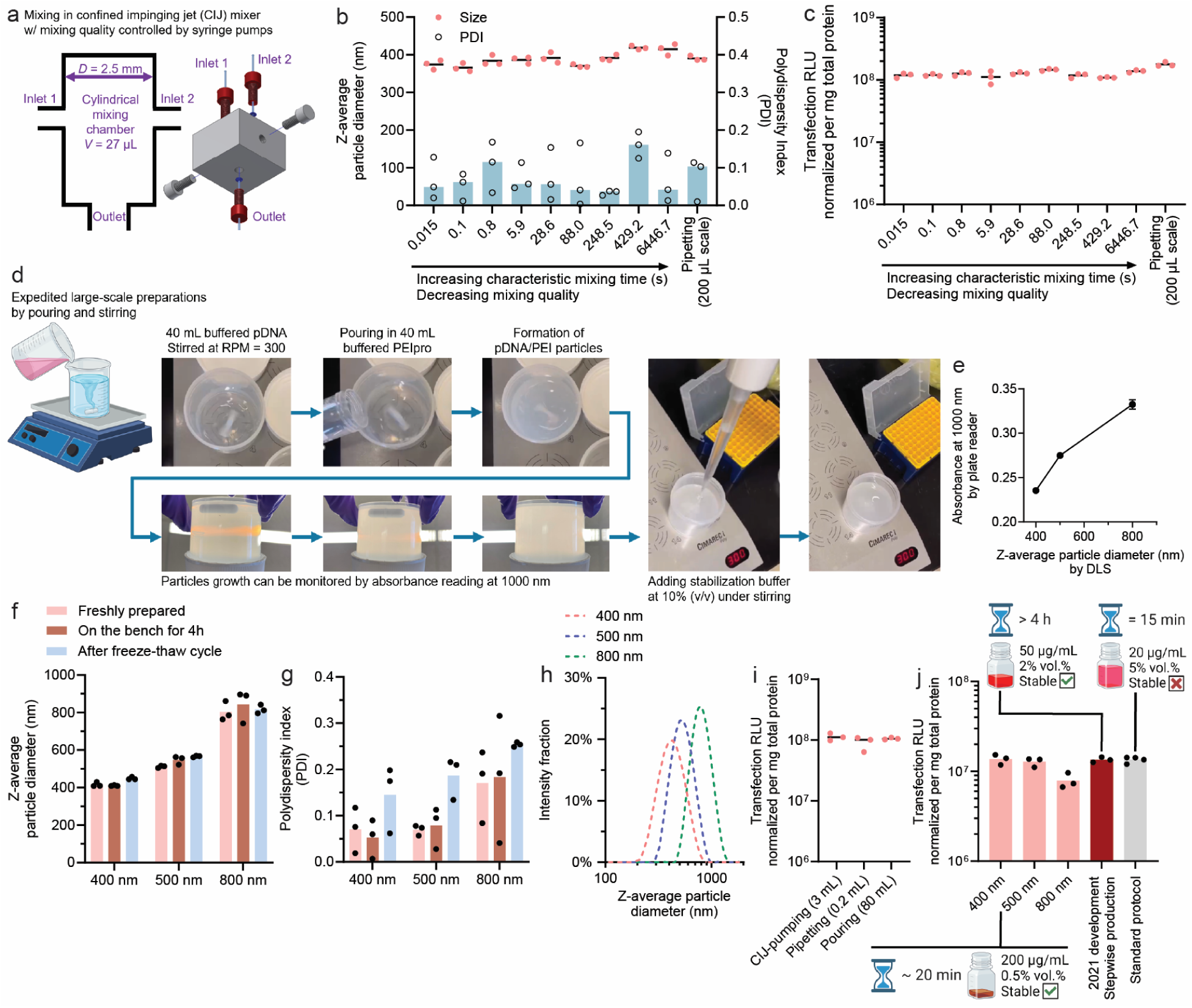
CBS-mediated assembly of stable, concentrated pDNA/PEI particles in different preparation scales. **(a)** The confined impinging jet (CIJ) mixer (coupled with syringe pumps) was used to vary the flow rate and characteristic mixing time (or mixing quality) of the buffered pDNA solution and the buffered PEIpro solution. The size and PDI **(b)**, and transfection efficiency **(c)** of 400-nm particles loaded with 10% (w/w) luciferase plasmid produced with different mixing quality, as controlled by the CIJ mixer or performed directly by pipetting. **(d)** The recordings of a particle production process executed at a scale of 80 mL and a pDNA concentration of 200 μg/mL, using easy-to-access labware under magnetic stirring and manual mixing. **(e)** Size growth monitored by measuring the absorbance of 100 μL of growing particles in a 96-well plate by a plate reader, which completes within seconds. The on-bench (4 h) and shelf (after a freeze-thaw cycle) stability of the particles produced at different sizes with an 80-mL scale in terms of the particle size **(f)** or the PDI **(g)** as assessed by DLS. **(h)** The size distribution of the particles produced on an 80-mL scale. **(i)** The transfection efficiency of 400-nm particles produced on an 80-mL scale compared to 400-nm particles prepared using the CIJ mixer or by pipetting. **(j)** The transfection of pDNA/PEIpro particles produced at different sizes with an 80-mL scale compared to 400-nm particles produced by our previously developed stepwise method or compared to particles prepared according to the manufacturer’s protocol for PEIpro. The insets specify the time needed to prepare the particles, the pDNA concentration, the relative culture volume ratio, and stability.

To fully demonstrate the wider applicability of this assembly method and scalability, we prepared pDNA/PEIpro at a batch size of 80 mL and selected a simple pouring method to mix 40 mL of PEIpro solution with 40 mL of pDNA solution in a 250-mL polypropylene (PP) beaker using a magnetic stirrer operating at 300 rpm (**Fig. 4d**). This mixing method is far less controlled than a typical pump-operated mixing condition. We also monitored particle size growth by measuring the UV absorbance at 1000 nm using a plate reader (**Fig. 4e**). Using a working curve reliably correlating DLS size of the particle suspension, linearly proportional to the absorbance at 1000 nm, we prepared stabilized particles at different target average sizes by pipetting the stabilization buffer into the suspension (**Fig. 4d**). Using such set-up, we generated three batches of particles with an average size of 400, 500 nm, and 800 nm, respectively. We verified their stability on the bench and after a freeze-thaw cycle by DLS (**Fig. 4f**). Low PDI values of around 0.1 were confirmed for the 400-nm and 500-nm particles (**Fig. 4g**). Combined with the unimodal size distribution of these particles (**Fig. 4h**), these results suggest that a high degree of uniformity was achieved with this crude scale-up mixing setup. The freeze-thaw process seemed to slightly increase PDI to around 0.2 (**Fig. 4g**). More importantly, the particles produced at this scale showed consistent transfection efficiency using a luciferase reporter pDNA in HEK293T cells, compared with particles generated at smaller scales by either pipetting or highly efficient CIJ mixing (**Fig. 4i**). At this production scale, our previously developed stepwise particle assembly method^22^ would take over 4 h to complete the production. The preparation protocol described here using the CBS buffer shortened the total assembly time to 15–45 minutes, which is substantially favorable.

### Transfection efficiency of the CBS-assembled shelf-stable pDNA/PEI particles

To benchmark our method of producing highly concentrated pDNA/PEIpro particles, we compared the transfection efficiency of CBS-assembled particles at 200 μg/mL with those produced using two industry-standard methods: (1) our previously published stepwise protocol, which generates pDNA/PEIpro particles at 50 μg/mL that was tested in LVV production^21^, and (2) the manufacturer’s PEIpro protocol, which generates non-stable complexes at 20 μg/mL. Despite the differences in preparation and concentration, the transfection efficiencies were comparable across all three formulations (**Fig. 4j**). This demonstrates that our method preserves functional performance, while offering distinct advantages of high operational stability and shelf stability, high pDNA concentration (up to 200 μg/mL), and a shorter preparation time of ∼20 min.

We then evaluated the size-dependent transfection efficiency of CBS-assembled particles in HEK293T (adherent culture) and HEK293F (suspension culture) cells using luciferase and tdTomato reporter plasmids. The transfection efficiency measured by the total luciferase expression level at 24 h (**Fig. 5a, b**), percentage of tdTomato^+^ cells characterized by flow cytometry (**Fig. 5c, d**), and average gene expression level per cell by the median fluorescent intensity (MFI) of tdTomato^+^ cells (**Fig. 5e, f**) revealed that the peak transfection efficiency were mediated by pDNA/PEI particles with an average size of 400 to 500 nm, showing the size-dependent profile consistent with our prior findings^21^.

**Figure 5.**
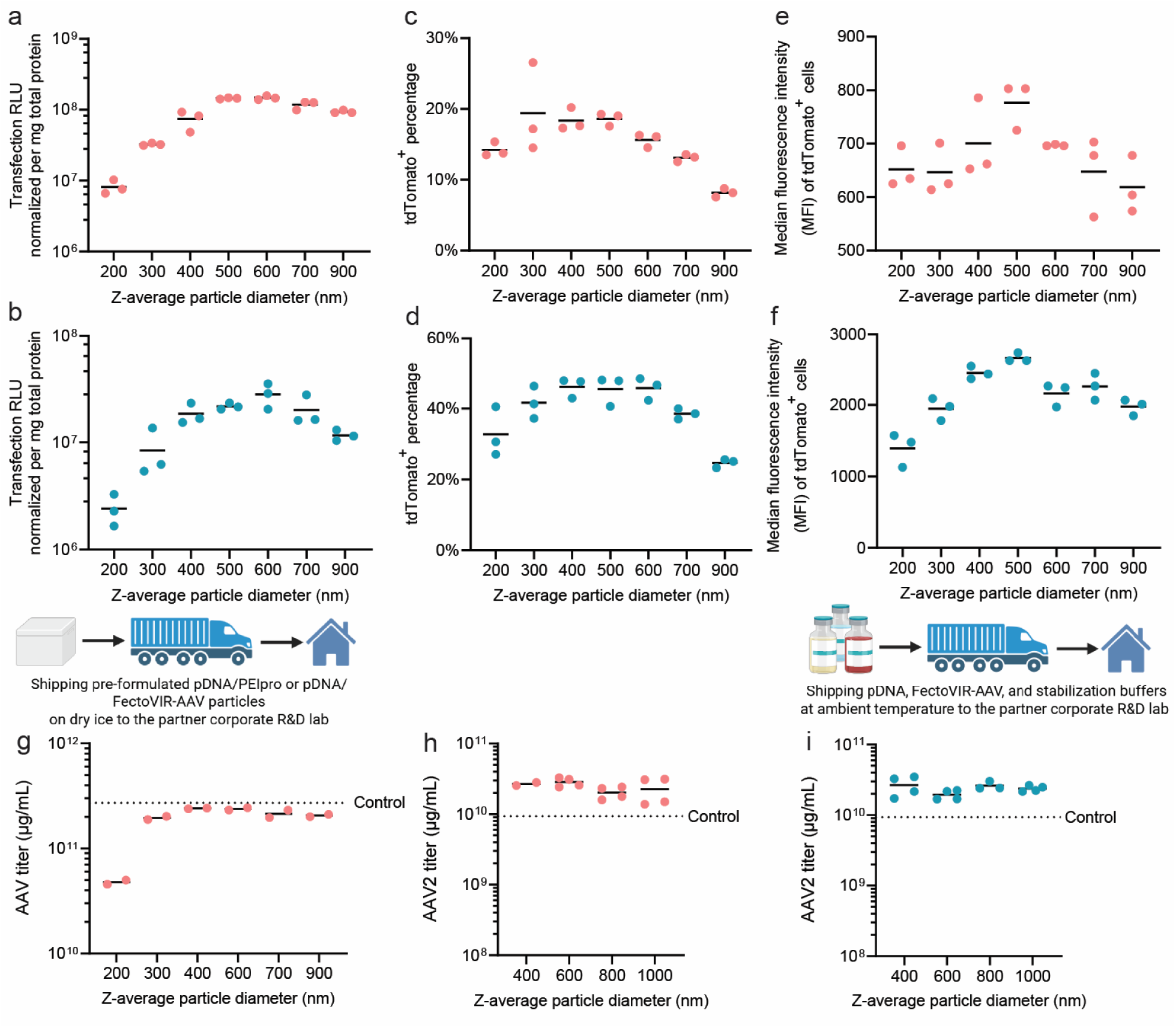
Transfection efficiency and rAAV production using CBS-assembled, concentrated and stable pDNA/PEI particles. (**a, b**) Transfection efficiency by different-sized particles loaded with 10% (w/w) luciferase plasmid, assessed at 24 h after transfection of HEK293T cells **(a)**, or at 48 h after transfection of HEK293F cells **(b)**. **(c, d)** Transfection efficiency by different-sized particles loaded with 100% tdTomato plasmid in terms of rate of transfection (tdTomato^+^), assessed at 24 h after transfection of HEK293T cells **(c)**, or at 48 h after transfection of HEK293F cells **(d)**. **(e, f)** Transfection efficiency of different-sized particles loaded with 100% tdTomato plasmid in terms of expression level per cell, *i.e.*, median fluorescent intensity or MFI of tdTomato^+^ cells, assessed at 24 h after transfection of HEK293T cells **(e)**, or at 48 h after transfection of HEK293F cells **(f)**. **(g)** rAAV titer from bioreactor production at Biogen Inc., transfected with pre-formulated stable pDNA/PEIpro particles at different sizes that were shipped on dry ice, thawed at ambient temperature for 2 h, and tested. **(h)** rAAV2 titer from bioreactor production at Polyplus Sartorius, transfected with preformulated pDNA/FectoVIR-AAV particles at different sizes that were shipped on dry ice, thawed at ambient temperature for 2 h, and tested. **(i)** rAAV2 titer from bioreactor production, transfected with on-site prepared, concentrated particles using the same set of buffers as in the experiment described in **(h)**, was shipped to and used by Polyplus Sartorius, as an independent verification of the method.

### AAV production using CBS-assembled shelf-stable transfection particles

To assess translational relevance, we collaborated with Biogen Inc. and Polyplus Sartorius to test our method in industrial AAV production settings. First, we used a plasmid cocktail developed by Biogen Inc. (USA, NASDAQ: BIIB) and formulated pDNA/PEIpro particles with an average size ranging from 200 to 900 nm in our Johns Hopkins University laboratory in Baltimore MD, using the CBS assembly protocol. The assembled particles were stored at –80°C before shipment on dry ice to the Biogen laboratory in Cambridge, MA, where the particles were stored at –80°C until use. On the day of experiments, the particles were thawed at ambient temperature for 2 h and then added to a benchtop bioreactor according to Biogen’s standard AAV production protocol. These particles with an average size of 300 to 900 nm yielded a similar AAV titer that matched the control level (**Fig. 5g**), which represents a typical titer obtained from the optimized Biogen protocol. This results proved the functionality, shelf stability, and bench stability of the pDNA/PEIpro particles assembled by CBS.

We further tested this particle assembly method using a new transfection reagent, FectoVIR-AAV, produced by Polyplus Sartorius. Citrate buffer was added to the solution set, and the pDNA/FectoVIR-AAV transfection particles were assembled using pre-screened conditions. The pDNA/FectoVIR-AAV particles were assembled at a batch size of 1.2 mL according to the above-described method. The stability of these particles flowing a freeze-thaw cycle was also confirmed (**Supplementary Fig. 4**). Particles with average sizes of 400, 600 nm, 800 nm, and 1000 nm were shipped to the Polyplus Sartorius laboratory in Illkirch, France and stored at –80°C until use. On the day of experiments, the particles were thawed at room temperature for 2 h and used for AAV2 production in benchtop bioreactors. These CBS-assembled particles showed comparable or higher AAV2 titers compared to pDNA/FectoVIR particles freshly formulated per the standard protocol (**Fig. 5h**). As a further validation, we sent buffer kits for on-site particle assembly at Polyplus Sartorius; the resulting AAV yields were highly consistent with those generated at Johns Hopkins, demonstrating reproducibility of the method (**Fig. 5i**).

## Conclusion

We report a simplified and scalable method for assembling shelf-stable, high-concentration pDNA/PEI particles, enabled by kinetic control of electrostatic complexation using trivalent citrate ions. This approach overcomes major barriers in transfection particle manufacturing, including reliance on precise mixing, low pDNA concentrations, and limited shelf life. The use of trivalent citrate anions reversibly reduces the positive charge density on PEI molecules, thereby reducing the complexation rate between pDNA and PEI at high concentrations and permitting fast production of pDNA/PEIpro and pDNA/FectoVIR-AAV transfection particles in the size range of 400 to 1000 nm with a high degree of uniformity. Our findings also demonstrate the unique roles of multi-valent ions in controlling electrostatic assembly of nanoparticles and opens up new engineering space of buffer valency for manufacturing transfection particles.

Using the citrate buffer saline developed, we consistently produced 400–500 nm pDNA/PEI particles within 20–40 minutes at 200 μg/mL, 10 to 20 times the typical concentration used in industry. This allowed a substantial reduction in the bioreactor dosing volume of the transfection particles. The CBS-assembled particles also show a high degree of on-bench stability for at least 4 h at ambient temperature, good shelf stability under cold storage at –80°C, and freeze-thaw stability. More importantly, this CBS-mediated assembly method is agnostic to the mixing method and batch scale. This platform may simplify liquid handling logistics, reduce process variability, and enhance manufacturing flexibility, making it suitable for wide adoption in viral vector production at various batch size scales. These features are particularly favorable for performing transfection in large-scale AAV production to develop more robust processes, and to potentially reduce the cost for AAV manufacturing. Beyond AAV production, this nanoparticle assembly protocol may have broad utility in enhancing the process robustness and efficiency of viral vector production at industry scales.

## Methods

### Preparation of stable, high-concentration pDNA/PEI particles

Plasmid DNA (gWiz-Luc, 6732 bp or tdTomato plasmid; both from Aldevron) was dissolved at 440 μg/mL in a pDNA dilution buffer containing 19% (w/w) trehalose. Poly(ethyleneimine) (PEIpro, Polyplus Sartorius, France) was similarly dissolved at 580.8 μg/mL in a PEI dilution buffer with 19% (w/w) trehalose. For kinetic control studies shown in **Figs. 2–4**, the dilution buffers included either phosphate-buffered saline (PBS; 137 mM NaCl, 2.7 mM KCl, 10 mM Na₂HPO₄, 1.8 mM KH₂PO₄) or citrate-buffered saline (CBS; 140 mM NaCl, 4 mM trisodium citrate, pH 7.0). For rAAV production, the buffers were optimized for pDNA cocktails from the corporate partners and for the FectoVIR carrier, which is proprietary to the corporate partners. The pDNA and PEI solutions were mixed at a 1:1 volume ratio, and the concentration of PEI was adjusted to achieve a nitrogen-to-phosphate (N/P) ratio of 5.5. When particles reached the target size monitored by optical density, a stabilization buffer containing trehalose and an acidic pH was added at a 10% v/v ratio to stop further size growth. Final particle suspensions contained 200 μg/mL pDNA and were either used immediately or stored at –80 °C.

### Control of mixing quality using a confined impinging jet (CIJ) mixer

To study the effect of mixing kinetics on particle assembly, we used a CIJ mixer that we previously developed^22, 29^. The PEI and pDNA solutions were loaded into separate syringes and injected into the CIJ device at controlled flow rates using a digital syringe pump (NE-4000, New Era Pump Systems). Flow rate determined the characteristic mixing time (τ_m_), allowing systematic evaluation of mixing effects on particle size and uniformity.

### Benchtop-scale particle production

For scale-up validation experiments described in **Fig. 4d–j**, 40 mL of pDNA solution was combined with 40 mL of PEI solution in a 250-mL polypropylene beaker under constant stirring at 300 rpm using a standard lab stirrer. Particle growth was monitored by measuring absorbance at 1000 nm using a 96-well plate reader. A calibration curve was established to correlate absorbance with DLS-derived particle size. Once particles reached the desired size (e.g., 400, 500, or 800 nm), stabilization buffer was added as described above.

### Characterization of the assembled pDNA/PEI particles

During complexation or after stabilization, the size of the particles was characterized by dynamic light scattering (DLS) using a Zetasizer ZS90 (Malvern, USA) at a 90° scattering angle. Z-average diameters (*D*_Z_) were reported throughout the study. The polydispersity index (PDI) was directly given by the instrument and is defined as *PDI* = (Δ⁄*D*_z_)^2^, in which Δ is the standard deviation of the size distribution. For morphology analysis, particles were deposited on glow-discharged lacey carbon TEM grids, negatively stained with 2% uranyl acetate, air-dried for 24 h, and imaged using a Talos F200C transmission electron microscope (Thermo Fisher Scientific).

### Cell culture and transfection experiments

For monolayer culture studies, HEK293T cells (American Type Culture Collection, ATCC, USA, maintained in DMEM supplemented by 10% FBS and 2 mM L-glutamine, at 37 ℃, 5% CO2, and saturated humidity) were seeded at a cell density of 100,000 cells/well in a 24-well plate, 1 day prior to transfection. Transfection particles at a DNA concentration of 200 μg/mL containing 10% (w/w) luciferase plasmid with 90% (w/w) non-coding plasmid or containing 100% tdTomato plasmid were diluted in Opti-MEM medium. The medium in each well of the plate was then replaced by particle-containing Opti-MEM. The stabilized particles were dosed into the cells at 1 μg DNA per well, and incubated with the cells for 4 h, followed by medium exchange with fresh cell culture medium and a subsequent culture of 44 h. Cells were then lysed with 200 μL reporter lysis buffer (Cat. #E4030, Promega Co.), and the lysates underwent a freeze-thaw cycle (–80 °C to ambient temperature). For luciferase measurements, 20 μL of each sample was mixed with 100 μL luciferase substrate (Cat. #E1483, Promega Co.) and the luminescence signal was measured by a luminometer. For tdTomato fluorescence assessment, the cells were detached by Trypsin-EDTA and stained with LIVE/DEAD™ Fixable Aqua Dead Cell Stain Kit (Thermo Fisher Scientific Inc.) for viability assessment before being analyzed by an Attune™ NxT Flow Cytometer (Thermo Fisher Scientific Inc.).

For suspension culture studies, HEK293F cells (Thermo Fisher Scientific, USA; maintained in FreeStyle 293 medium, at 37 ℃, 8% CO2, and saturated humidity) were seeded at 1,000,000 cells/well in a 12-well plate, on the day of transfection. Stabilized particles at a DNA concentration of 200 μg/mL containing 10% luciferase plasmid with 90% non-coding plasmid, or 100% tdTomato plasmid, were added directly into the cell suspension to reach a dosage of 1 μg DNA per well. Cells were incubated with the particles for 48 h until analysis. For luciferase assessment, cells were collected from the plates, pelleted by centrifugation at 300 ×g for 5 min, and then lysed with 500 μL reporter lysis buffer. The luminescence assay was carried out using the same protocol as described above; The tdTomato fluorescence assessment using flow cytometry was conducted using the same protocol as described above.

### Molecular dynamics (MD) simulations

MD simulations presented in this manuscript were performed in GROMCAS^32^ with Martini force field parameters^29, 33, 34^. These coarse-grained simulations allowed us to run for longer and relatively large systems, which would not have been possible with atomistic simulations. We used Salassi et al.^33^ for modeling citrate ion and Mahajan and Tang^34^ parameters for PEI and 40% of the monomers were charged. Fifty small PEIs (14-monomers) were randomly placed in a 20×20×20 nm^3^ simulation box with 140 mM NaCl and added 10 mM of phosphate or citrate ions. First, energy minimization was performed on the systems, followed by initial equilibration under NVT ensemble for 50 ns with 10 fs timesteps. The final equilibration simulation was performed for 5 μs with 20 fs timestep. Neighbor searching was performed up to a cut-off distance of 1.2 nm using the Verlet particle-based approach. The potential-shift method was applied for the short-range Lennard-Jones (LJ) 12-6 interactions at a cut-off of 1.2 nm. The reaction-field method was used for calculations of the non-bonded interactions between charge beads (cutoff 1.2 nm). The dielectric constant was fixed at 15 according to the Martini model. The bonds were constrained with the LINCS algorithm^35^. The temperature of the system was maintained using the v-rescale algorithm at a reference temperature of 298 K. The isotropic Parrinello-Rahman barostat was utilized with the reference pressure of 1 bar^36^. Ovito was used for clustering and visualizations^37^, and a cluster cutoff of 0.72 nm was selected based on the first valley of the Martini bead radial distribution function^29^. To measure the aggregation process, we calculated the number-average cluster size (CSN) as *CSN* = ∑ *N*· *M*⁄∑ *M*, where N is the cluster size, and M is the concentration of the clusters. If at least two monomers were within 0.72 nm of each other, we defined them as forming a cluster. Similar simulation procedures were followed for simulating large PEI, although only 1 PEI was placed in the simulation box (**Fig. 3e**).

For calculating the free energy of pDNA/PEI binding, we first placed a 24 bp DNA (Sequence: dCdGdCdGdAdAdTdTdCdGdCdGdCdGdCdGdAdAdTdTdCdGdCdG) in a simulation box and added a PEI near it. After 1 μs of equilibration, we pulled the PEI from the DNA in the radial direction using a constant force. From different distances between DNA and PEI, we selected reaction coordinates that were 1.5 Å apart. We performed umbrella sampling in those coordinates using a force constant of 1000 kJ/mol/nm^2^ and used the WHAM algorithm to get the binding free energy^38^. Each reaction coordinate was simulated for 700 ns, of which the last 500 ns of data were used during the WHAM algorithm.

The methods to perform the MD simulations shown in **Supplementary Fig. S3** are described in **Supplementary Methods.**

## Acknowledgements

This work was partially supported by sponsored research agreements with Biogen Inc. and Polyplus Sartorius, and by the National Institutes of Health under grant R01CA281143 awarded to H.-Q.M. The authors thank all collaborators from Biogen and Polyplus Sartorius for their partnership in validating the methods and sharing production insights. The computational work was carried out at the Advanced Research Computing at Hopkins (ARCH) core facility (rockfish.jhu.edu), which was supported by the National Science Foundation (NSF) grant OAC 1920103.

## Author Contributions

Y.H. and H.-Q.M. conceived and supervised the project. J.L., D.Y., J.M., and N.K. conducted experiments on particle assembly, characterization, and cell transfection. T.H.P. and T.C. performed molecular dynamics simulations and contributed to the mechanistic analysis. P.B. and M.C. carried out AAV production experiments and data analysis at Biogen. M.G. conducted experiments at Polyplus Sartorius and analyzed AAV titers. S.L., Y.Z., L.C., and T.-H.W. contributed to data interpretation and figure generation. J.L., Y.H., and H.-Q.M. wrote the manuscript with input from all co-authors.

## Competing Interests

J.L., Y.H., and H.-Q.M. are co-inventors on a patent application covering the pDNA/PEI particle assembly technique described in this study, filed through and managed by the Johns Hopkins Office of Technology Ventures. P.B. and M.C. are employees of Biogen; M.G. is an employee of Polyplus Sartorius. The other authors declare no competing interests.

