## Supplementary Information for "Buffer Valency Engineering Enables High-concentration and Shelf-stable DNA Transfection Particles for Viral Vector Production"

[1]Institute for NanoBioTechnology, Johns Hopkins University, Baltimore, MD, USA. [2]Department of Materials Science and Engineering, Johns Hopkins University, Baltimore, MD, USA. [3]Translational Tissue Engineering Center, Johns Hopkins University School of Medicine, Baltimore, MD, USA. [4]Department of Biomedical Engineering, Johns Hopkins University School of Medicine, Baltimore, MD, USA. [5]Polyplus Sartorius, Illkirch-Graffenstaden, France. [6]Biogen Inc., Cambridge, MA, USA. [7]Department of Mechanical Engineering, Johns Hopkins University, Baltimore, MD, USA. [8]Department of Chemical and Biomolecular Engineering, Johns Hopkins University, Baltimore, MD, USA. [9]Present address: David H. Koch Institute for Integrative Cancer Research, Massachusetts Institute of Technology, Cambridge, MA, USA.

[10]These authors contributed equally to this work.

**Supplementary Methods**

MD simulations presented in this Supplementary Information are performed in LAMMPS molecular dynamics package with reduced Lennard-Jones (LJ) units. The system consists of PEI, DNA, and ions (phosphate and citrate), modeled using a coarse-grained bead-spring representation. Non-bonded interactions are modeled using the Weeks–Chandler–Andersen (WCA) potential:

$$V(r,b) = \begin{cases} 4\epsilon_{LJ}[(b/r)^{12} - (b/r)^{6} + 1/4], & r \le r_c \\ 0, & r > r_c \end{cases}$$

We used $\sigma = 0.32$ nm as the unit of length and $\epsilon_{LJ} = 1$ sets the energy scale. $b_{DNA,DNA} = 3.12\sigma$, $b_{DNA,PEI} = 3.12\sigma$ and all other bead pairs: $b = \sigma$. The cutoff for all interactions is fixed at $r_c = 2^{1/6}b$, ensuring purely repulsive interactions. To capture the intrinsic rigidity of DNA and PEI, angular potential was applied according to respective persistence length. Electrostatic interactions were computed using the Debye–Hückel approximation, with a screening length corresponding to an ionic strength of ~100 mM. Additional multivalent ions (10 mM concentration) were modeled explicitly: divalent ions as single charged beads, and citrate ions as linear trimers of three charged beads connected by stiff harmonic bonds, capturing their trivalent, multidentate nature. Simulation time step was 0.002. All simulations used the velocity-Verlet integrator with a Langevin thermostat. Periodic boundary conditions were applied in all directions.

**For supplementary Fig. 3a.** The system contained a 300 bp DNA chain and PEI chains with 80 monomers each. Initial equilibration was performed in the absence of attractive DNA–PEI interactions to allow PEI complexation with ions (5,000,000 steps). This mimics pre-complexation in solution. Attractive DNA–PEI interactions were then introduced to study complex formation and morphology (10,000,000 steps).

**For Supplementary Fig. 3b:** Citrate ions were modeled as linear trimers of three charged beads connected by stiff harmonic bonds. The PEI chains contained 80 monomers; Divalent ions were modeled as single charged beads. The PEI chains contained 80 monomers.

### Supplementary Figures

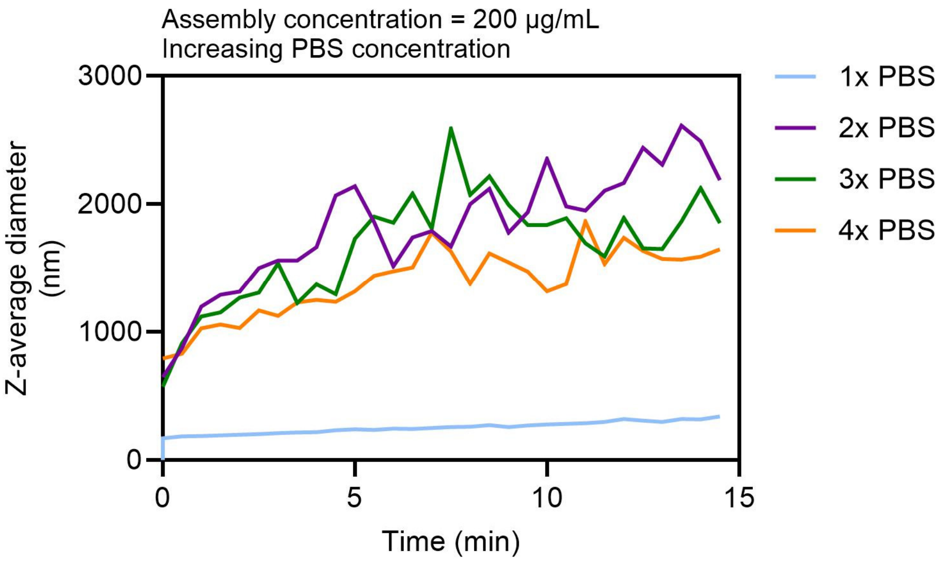


**Supplementary Fig. 1. Particle growth at a pDNA concentration of 200 µg/mL mediated by different concentrations of PBS as a buffer.**

While increasing the concentration of PBS enhanced charge screening by the increased ionic strength from monovalent and divalent ions, it did not achieve the steady, uniform particle assembly as mediated by citrate-buffered saline.

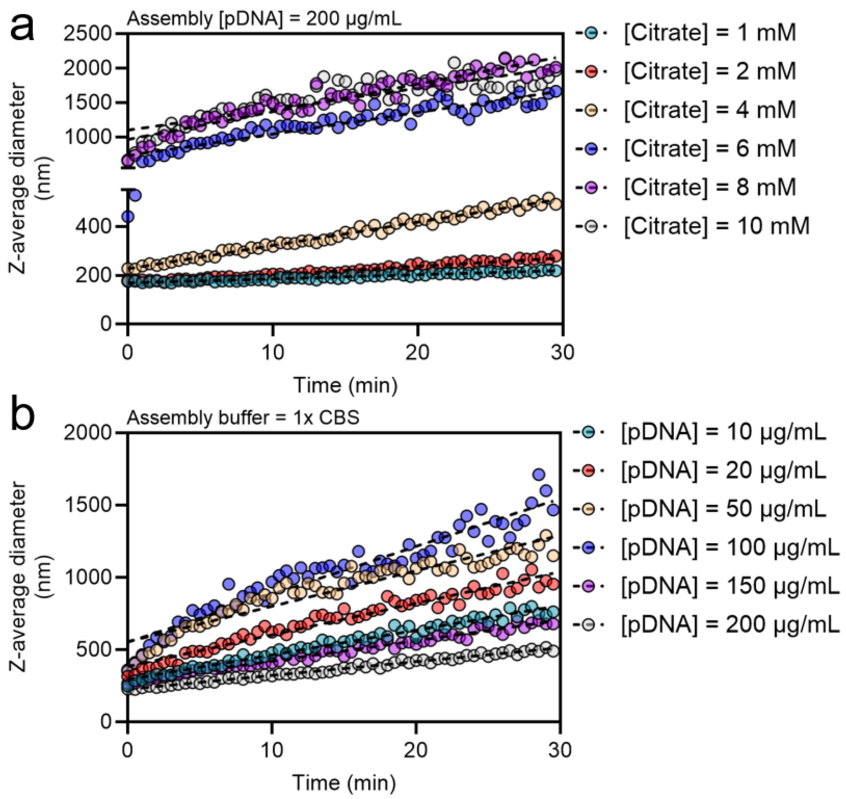


**Supplementary Fig. 2. NPa kinetics**

The effects of **(a)** plasmid DNA concentration [pDNA] and **(b)** citrate concentration [Citrate] added to the phosphate buffer on the nanoparticle assembly (NPa) process following polyelectrolyte complexation (PECn) step.

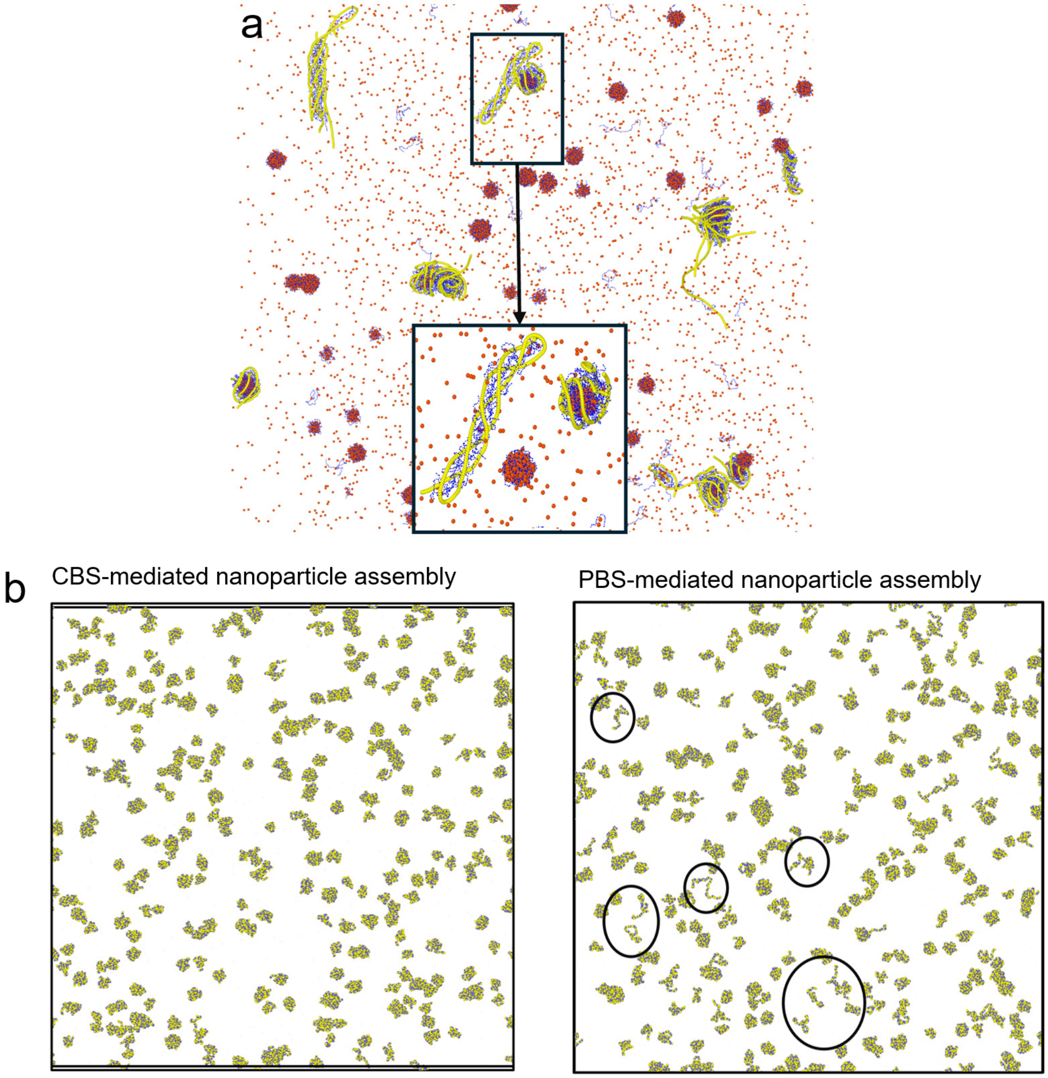


**Supplementary Fig. 3. Molecular dynamics simulations for nanoparticle assembly (NPa) step following the initial pDNA-PEI polyelectrolyte complexation (PECn) step.**

**(a)** PEI (blue), pDNA (yellow), and anions (red) were included as the three main components in the system. Over time, we observed the aggregation between anions and PEI molecules, and then some portions of anions were replaced by DNA molecules, which provide stronger binding with PEI due to a higher density of negative charges. **(b)** The incorporation of the trivalent anions of citrate in the PEC resulted in the formation of assembled particles with a more regular, spherical shape, which further accelerates the aggregation between PEC seeds as expected from the DLVO theory, eventually leading to submicron particles. In contrast, PBS-mediated assembly resulted in the formation of assembled particles with more irregular shapes. These simulation findings are consistent with TEM observations.

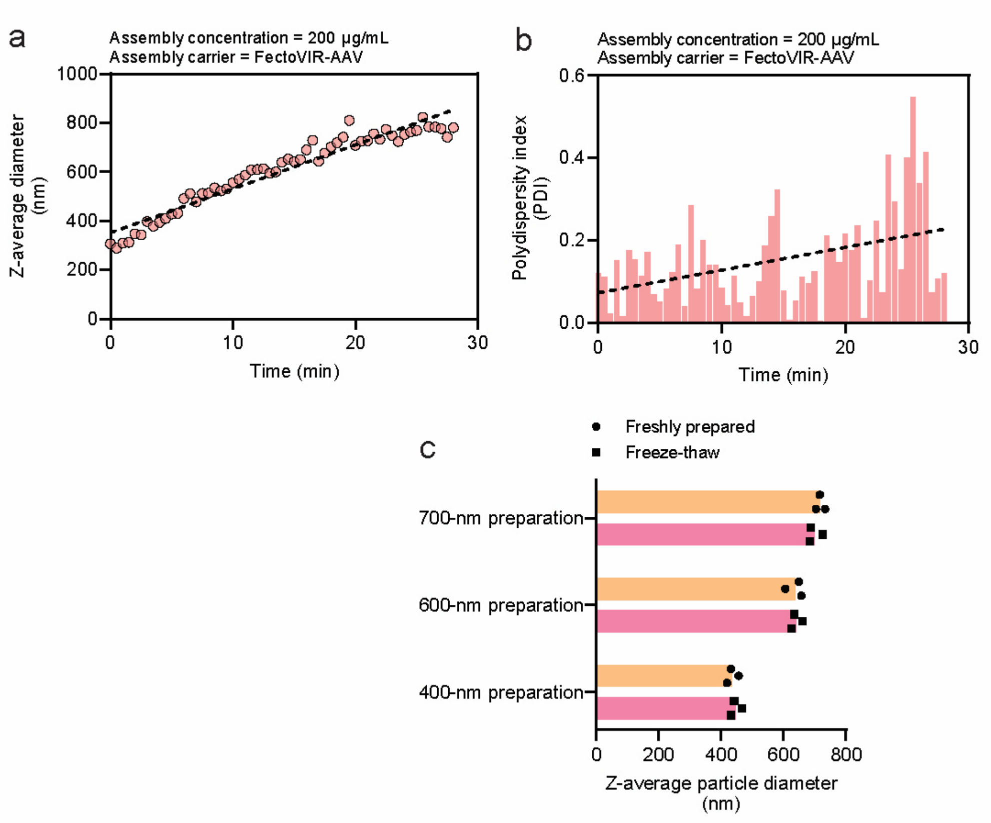


**Supplementary Fig. 4. Citrate-buffered saline (CBS)-mediated particle growth using Fecto-VIR AAV carrier.**

**(a)** Dynamic light scattering (DLS) size measurements during a particle assembly process mediated by CBS. **(b)** DLS polydispersity index (PDI) measurements during a particle assembly process mediated by CBS. **(c)** The freeze-thaw stability of particles assembled at different sizes by CBS.

**Supplementary Table 1.** Theoretical characteristic mixing time correlated with mixing rates.

| **Total flow rate in mixing (mL/min)** | **40** | **20** | **10** | **6** | **4** | **3** | **2.3** | **2** | **1** |
|---|---|---|---|---|---|---|---|---|---|
| **Theoretical characteristic mixing time (s)** | 0.015 | 0.1 | 0.8 | 5.9 | 28.6 | 88.0 | 248.5 | 429.2 | 6446.7 |
